# Involvement of birth weight and body composition on autonomic recovery following aerobic exercise in children: A prospective, observational and analytical study

**DOI:** 10.1101/521500

**Authors:** Juliana Edwiges Martinez Spada, Fernando R. Oliveira, David M. Garner, Vitor E. Valenti

**Affiliations:** Post-Graduate Program in Physical Therapy, UNESP, Presidente Prudente, SP, Brazil.; School of Public Health, University of Sao Paulo, Sao Paulo, SP, Brazil.; Cardiorespiratory Research Group, Department of Biological and Medical Sciences, Faculty of Health and Life Sciences, Oxford Brookes University, Headington, Gipsy Lane, Oxford, OX3 0BP, United Kingdom.

**Keywords:** Autonomic Nervous System, Birth weight, Children, Exercise, Heart Rate

## Abstract

Birth weight (BW) can be used to assess the health status of the newborn. However, its impacts on later in life regarding heart rate (HR) variability (HRV) is not totally clear. We aimed to analyze the involvement of BW and body composition on HRV recovery following aerobic exercise in children. The study was conducted in healthy children 9 to 11 years of age (40 females and 27 males) divided into two groups: G1 (BW < 3400 grams, N = 33) and G2 (BW > 3400 grams, N = 34). The volunteers completed an experimental protocol of submaximal aerobic exercise on a treadmill and remained seated for 30 minutes after exercise. Systolic (SAP) and diastolic arterial pressure (DAP), respiratory rate (f) and HRV were analyzed before and during recovery from exercise. SAP and f were significantly decreased 30 minutes after exercise compared to 1 minute after exercise in G1 and G2. Mean HR, high frequency band of spectral analysis (HF), root mean square of successive interbeat intervals difference, SD1 index and mean lenght were diminished 0 to 5 minutes after exercise compared to rest in G2 while maximum lenght increased 0 to 5 minutes after exercise compared to resting in G2. Linear regression revealed association of fat percentage and BW with nonlinear HRV recovery. In conclusion, autonomic recovery after exercise was somewhat delayed in children with high BW. BW and fat percentage *slightly* influence HRV recovery.

## INTRODUCTION

The cardiovascular system is regulated through the autonomic nervous system (ANS). Its sympathetic and parasympathetic components sustain the organism within its homeostatic patterns [1]. The ANS regulates the respiratory, thermoregulatory and vasomotor systems, baroceptors and endocrine metabolism (renin-angiotensin-aldosterone) [2]. Activation of the sympathetic nervous system elevates heart rate (HR), increases cardiac contractility, reduces venous compliance and induces vasoconstriction, whilst parasympathetic modulation lessens HR through vagal impulses [3].

The autonomic responses of HR during aerobic exercise are considered by vagal withdrawal in the first few seconds that elevate HR [4]. Afterwards, an increase in sympathetic activity increases cardiac contraction and accelerates wave ventricular depolarization conduction^2^. In contrast, non-elevation of HR in the initial phase of exercise may be indicative of a deficiency of vagal activity [3]. Vagal re-entrance and sympathetic withdrawal contribute to the return of HR levels attained at the start of exercise [4].

HR variability (HRV) evaluates interbeat oscillations [5,6] and can be noninvasively investigated through the analysis of RR intervals in physiological or pathological conditions, providing evidence regarding the influence of ANS [7,8] on the sinus node through linear and non-linear methods in the time and frequency domains by application of nonlinear analysis through complexity techniques or algorithms [9].

Autonomic recovery following exercise corresponds to the rate of HR decline owing to reactivation and coordinated vagal withdrawal following exercise [10–12]. Examination of autonomic recovery after aerobic exercise indicates appropriate clinical data regarding autonomic imbalance and is correlated to a reduction of vagal tone or exaggerated sympathetic activation [13,14]. Prior studies evaluating the autonomic recovery after exercise demonstrated that HR recovery after exercise is influenced by features related to the anthropometric variables and exercise characteristics [15–18].

Birth weight (BW) is a interesting and important anthropometric variable that merits attention. Battaglia and Lubchenco [19] developed an intrauterine growth chart for the classification of BW at gestational age. Furthermore, previous studies have revealed that adult individuals with low BW had autonomic dysfunction at rest, characterized by increased sympathetic activity and lessened parasympathetic activity [20–21]. Still, different outcomes regarding the relationship between low BW and autonomic dysfunction have demonstrated ambiguous and inconclusive results [22].

A previous study investigated 5 to 12 year-old children during sleep and, it was observed that the association between low BW and prematurity was connected to cardiac structure alterations; yet, it was unable to modify the autonomic control [23]. In a sample of 397 children, the parasympathetic modulation at rest was reduced in adults with low BW, and those with higher BW had greater parasympathetic and baroreflex activity, indicating that autonomic control can be modified in adults born with low BW [24].

Taken together, it leads us to hypothesize that children with higher BW would achieve faster autonomic recovery following exercise.

It should be highlighted that there are physiological and maturational idiosyncrasies between boys and girls of the same age that are evident in the pubertal period [25]. Nevertheless, Goto *et al*. [26] was unable to find modifications between the genders. Similarly, Guilkey *et al*. [27] found no discrepancies in HRV between boys and girls 9 to 11 years old after maximal and submaximal exercise. Of late, Souza *et al*. [22] reported no significant changes in HRV between boys and girls aged 5 to 14 years old. In this study, we decided to perform the investigation of data considering boys and girls in the same group.

After an extensive the literature review, we discovered no studies that evaluated the impact of BW on HRV during recovery from exercise in children with a restricted age (9 to 11 years). Bearing in mind that the hemodynamic response to exercise may provide evidence that is unable to be detected at rest [18]; we draw attention to the relevance of a study that emphasizes children who do not apparently present cardiorespiratory diseases. This would assist the identification of possible predispositions to pathological conditions. So, we investigated the involvement of BW and body composition on autonomic recovery after aerobic exercise in children.

## METHODS

### STROBE Guidelines

This investigation conforms to the STROBE (STrengthening the Reporting of OBservational studies in Epidemiology) guidelines. Our study contains details of the study design, participants, setting, measurement, variables, description of potential sources of bias, data sources, quantitative variables description, and statistical methods.

### Population study and Eligibility Criteria

This study was performed in 67 healthy term-born (BW > 2500 grams, gestational age > 39 weeks) children between 9 and 11 years old split into two groups according to the BW (3400 grams): G1 (BW < 3400 grams, 15 boys and 18 girls) and G2 (BW > 3400 grams, 12 boys and 22 girls). We chosed 3400 grams because it was the median of the entire group.

The prenatal evidence from the mother’s report was confirmed with her medical records. Family medical histories were attained using a questionnaire completed. We excluded children with impairments that circumvented the correct performance of the procedures (cardiovascular, renal, respiratory, endocrine, orthopedic and neurological disorders), those under pharmacological treatments that influence the ANS, or girls who had started their menstrual cycle.

### Ethical approval and informed consent

All study protocols were approved by the Research Ethics Committee in Research of UNESP/Marilia (CAAE – 75760117.4.0000.5406) and a statement explicitly stated that the methods were undertaken in accordance with the 466/2012 resolution of the National Health Council of 12/12/2012. Informed consent was attained from all children’s parents that signed a confidential consent letter.

### Study design and Setting

This is a prospective, observational and analytical sectional study performed at the Autonomic Nervous System Study Center at UNESP/Marilia, SP, Brazil.

### Bias

The introductory examination was completed to evaluate the eligibility criteria and to obtain descriptive statistical characterizations about the individuals. An anamnesis was made to confirm the absence of recognized diseases and treatment with medications. The descriptive profile of the subjects was defined to characterize the sample, reduce the unpredictability of the variables, improving reproducibility and physiological interpretation. Before the start of the experimental procedures, subjects were documented according to age, mass (kg), height (m), systolic (mmHg) and diastolic arterial pressure (mmHg), waist (WC), hip (HC) and abdominal (AC) circumferences, waist-to-hip ratio and body mass index (BMI).

The protocol was undertaken individually between 1:00 pm and 6:00 pm to standardize circadian influences on HRV [27]. It was also performed in a soundproofed room with controlled temperature between 22 °C and 25 °C and humidity amid 50% and 60% at the School of technical courses in informatics - Igeeks^®^ (Tupis, 236, 17.600-000, Tupã, SP, Brazil).

The children were told to avoid drinking beverages containing stimulants or caffeinated drinks for 24 hours prior to the evaluation, food 8 hours before the assessment. They were instructed not to perform strenuous exercises for 24 hours before appraisal. Appropriate and comfortable clothing should be worn to undergo the necessary physical exertions.

### Initial assessment and Experimental Protocols

In the initial assessment the researcher logged: age, gender, weight, height, gestational age, BW, systolic (SAP) and diastolic arterial pressure (DAP), HR and respiratory rate (f) and whether their parents presented with cardiovascular disease.

WC, HC and AC were attained in orthostatism, with the abdomen relaxed and arms extended along the body, being measured with a tape measure positioned in the area of lesser curvature located between the final rib and the iliac crest. Waist-stature (WSR) and waist-hip ratio (WHR) were considered. BMI was attained according to the recommendations described by Lohman *et al*. [28]. Body adiposity index (BAI) [29] and conicity index (CI) [30] were gained through anthropometric data, and the body fat percentage (BF%) was estimated via bioimpedance [31,32].

HR was recorded with the Polar RS800cx HR monitor (Polar Electro, Finland) and respiratory rate (f) was enumerated by counting the respiratory cycles during one minute whilst the volunteer was uninformed about the procedure taking place. Thus, avoiding influences and consequent changes in the subjects’ f.

SAP and DAP were measured indirectly by auscultation using a calibrated aneroid sphygmomanometer and stethoscope (Premium, Barueri, São Paulo, Brazil) on the left arm whilst the individual continued seated and breathing spontaneously.

To avoid measurement distortions, the same researcher measured the same parameters throughout the whole experiment.

After the preliminary evaluation, the HR monitor was located on the subjects’ thorax, at the level of the distal third of the sternum. Before recording HR, SAP and DAP were measured.

The children remained at rest for 15 minutes in the seated position under spontaneous breathing, followed by 5 initial minutes walking on a treadmill for physical ‘warming up’ (50 - 55% of maximal HR (HRmax) HR: 208 – 0.7 × age) [34], next 25 minutes with 60-65% without inclination, and increments of 0.5 km/h every minute until reaching submaximal HR.

Directly after the exercise, subjects underwent one minute standing and then subsequently seated for passive recovery for a further 29 continuous minutes, totaling 30 minutes of recovery. During recovery from exercise volunteers remained seated in silence with spontaneous breathing, they did not sleep, did not perform any movements that would induce autonomic changes and did not ingest any type of drink or food.

HR, f, SAP and DAP were logged at 15 minutes of rest and at 1 and 30 minutes during recovery from exercise. HRV analysis was achieved at the following times: Rest (10^th^ to 15^th^ minute of resting) and during recovery: 0 to 5^th^ minute, 5^th^ to10^th^ minute, 10^th^ to 15^th^ minute, 15^th^ to 20^th^ minute, 20^th^ to 25^th^ minute and 25^th^to 30^th^ minute [35].

### Variables, Data Sources and Outcome Measures

#### HRV analysis

RR intervals were recorded by the cardiac portable monitor. The datasets were transferred to the Polar Pro Trainer program (v.3.0, Polar Electro, Finland). Digital filtering followed by manual filtering was accomplished for the disposal of artifacts. For data analysis, RR intervals were selected for analysis and extracted into a ‘txt’ file. All indices were evaluated using a fixed number of 256 consecutive stable RR intervals obtained from the baseline ending as well as the final 256 intervals of each recovery period. Only series with greater than 95% of sinus beats were included in the study.

HRV analysis followed directives from the Task Force^36^ and have been previously published [37,38]. Kubios HRV^®^ software (Kubios^®^ HRV v.1.1 for Windows, Biomedical Signal Analysis Group, Department of Applied Physics, University of Kuopio, Finland) was necessary to calculate linear indices.

The time domain index was analyzed via the rMSSD indices (square root of the mean of the square of the differences between adjacent normal RR intervals in a time interval) expressed in milliseconds (ms) and SD1 (instantaneous variability of beat-to-beating). For the frequency domain analysis, the high frequency spectral component (HF) calculated through the Fast Fourier Transform was used in absolute units (ms^2^) [39].

The symbolic analysis of HRV was completed through the distribution of the series of RR intervals at the levels: 0V and 2ULV through CardioSeries v2.4^®^ software (Ribeirao Preto, SP, Brazil). Detailed information concerning symbolic analysis has been described previously [40].

The reccurence analysis of HRV was achieved quantitatively using the Recurrence Plot (RP) through the Visual Recurrence Analysis^®^ software to investigate the following indices: mean lenght (L Mean), recurrence rate (REC), determinism (DET), laminarity (LAM), Shannon Entropy (Shannon Shannon Entropy, SE) and maximum lenght (L Max).

#### Study Size

The sample size was computed using a pilot test, wherein the online software provided by the website www.lee.dante.br was necessary taking into consideration the RMSSD index as a variable. The significant difference in magnitude assumed was 14.11 ms, with a standard deviation of 12.8 ms, per alpha risk of 5% and beta of 80%. The sample size designated a minimum of 13 individuals per group.

#### Statistical analysis

Data was presented as descriptive statistics to characterize the sample and were designated by the statistical values of mean, standard deviation and 95% confidence intervals.

Data normality was assessed via the Shapiro-Wilk test. The unpaired Student t test (parametric) or the Mann-Whitney test (non-parametric) were obligatory to compare descriptive characteristics between the groups.

For comparison between the moments (rest vs. recovery from exercise), the repeated measurements one-way analysis of variance (ANOVA1) test followed by Dunnett’s test (parametric distribution) or the Friedman test followed by Dunn’s test (non-parametric distribution) were required. The two-way repeated measures analysis of variance technique (ANOVA2) was performed to analyze any differences between groups (birth weight) vs. time (recovery time points). The data of the repeated measurements were checked for sphericity violation using the Mauchly test and the Greenhouse-Geisser correction was applied when the sphericity was violated.

Significant differences were considered statistically significant when the p-value was lower than 0.05 (or <5%). The statistical analysis were performed with Minitab software (Minitab, PA, USA) and GraphPad InStat - v3.06 (GraphPad Software, Inc., San Diego California USA).

Effect size was calculated through Cohen’s *d*. Large effect size was considered for values > 0.9 and medium effect size for values between 0.9 and 0.5 [41].

We performed correlation of HRV with BW, body adiposity index, fat percentage through Spearman correlation coefficient. We considered high correlation for r > 0.75 and moderate correlation for r between 0.75 and 0.5. We applied simple linear regression models to model 0V, Recurrent, Determinism - DET (%), Percent Laminarity and Shannon Shannon Entropy parameters as dependent variables and fat percentage and BW as independent variables. As a result of the non-normality of Shannon Shannon Entropy to fit the regression model through the cubic method prior to the analysis Determinism - DET and Percent Laminarity it was not possible to complete the transformation to normality, so the stated parameters were excluded from the regression analysis.

## RESULTS

We acknowledged that 28% of children’s parents in G1 exhibited cardiovascular impairment and 21% in G2 reported cardiovascular disease.

According to Table 1, children in the G2 had a higher gestational age, BW and height (p<0.05) whilst children in the G1 group had elevated body adiposity index and fat percentage (Table 1).

**Table 1:** Descriptive statistics of age, gestational age, weight, birth weight, height, waist (WC), hip (HC) and abdominal (AC) circumference, waist-hip circumference, waist-stature circumference, body mass index, conicity index, dody adiposity index and fat percentage.

| Variables | G1 | G2 | P | Cohen |
| --- | --- | --- | --- | --- |
| **Age (years)** | 9.62±0.70<br>[9.37-9.88] | 9.72±0.62<br>[9.50-9.94] | 0.57 | - |
| **Gestational age (weeks)** | 37.90±1.71<br>[37.2-38.5] | 39.48±0.93<br>[39.1-38.8] | < 0.0001**** | 0.1 |
| **Weight (Kg)** | 39.84±8.87<br>[36.6-43.0] | 39.16±7.69<br>[36.4-41.8] | 0.59 | - |
| **Birth weight (Kg)** | 2.88±0.34<br>[2.76-3.00] | 3.74±0.39<br>[3.60-3.88] | < 0.0001**** | 2.3 |
| **Height (m)** | 1.39±0.06<br>[1.37-1.42] | 1.43±0.08<br>[1.40-1.46] | <0.05* | 0.5 |
| **Waist circumference (cm)** | 67.8±9.56<br>[64.3-71.2] | 65.6±8.95<br>[62.4-68.8] | 0.33 | - |
| **Waist-hip ratio (cm)** | 81.31±7.57<br>[78.58-84.04] | 79.51±7.96<br>[76.68-82.34] | 0.35 | - |
| **Abdominal circumference (cm)** | 74.18±10.31<br>[70.47-77.90] | 69.51±9.48<br>[66.15-72.87] | 0.06 | - |
| **Waist-stature ratio (cm)** | 0.48±0.05<br>[0.46-0.50] | 0.45±0.05<br>[0.43-0.47] | 0.04 | 0.6 |
| **Waist-hip ratio (cm)** | 0.83±0.04<br>[0.81-0.84] | 0.83±0.13<br>[0.78-0.87] | 0.09 | - |
| **Body mass index (Kg/m<sup>2</sup>)** | 20.24±3.28<br>[19.05-21.42] | 18.82±2.77<br>[17.83-19.81] | 0.06 | - |
| **Conicity index** | 1.16±0.06<br>[1.14-1.19] | 1.15±0.10<br>[1.12-1.19] | 0.27 | - |
| **Body adiposity index** | 31.26±3.51 | 28.25±4.92 | 0.01** | 0.7 |
|  | [29.99-32.53] | [26.51-30.00] |  |  |
| **Fat percentage (%)** | 25.81±7.60 | 20.27±9.03 |  |  |
|  | [22.81-28.82] | [16.77-23-78] | 0.01** | 0.6 |

Mean ± standard deviation [95% Confidence interval].
**Legend:** G1: birth weight < 3400 grams; G2: birth weight > 3400 grams; Kg: kilogram; m: meters; cm: centimeters; %: percentage. \* p<0.05; \*\* p<0.01; \*\*\* p<0.001; \*\*\*\* p<0.0001.

SAP was significantly reduced 1 minute (Cohen’s *d*: 0.03) and 30 minutes (Cohen’s *d*: 1.05, large effect size) after exercise compared to rest in G1. In G2 SAP was also decreased 1 minute (Cohen’s *d*: 0.39, large effect size) and 30 minutes (Cohen’s *d*: 1.04, large effect size) following exercise. We also recognized that f declined 1 minute (Cohen’s *d*: 0.23, small effect size) and 30 minutes after exercise (Cohen’s *d*: 1.22, large effect size) in G1. In accordance, f was decreased 1 minute (Cohen’s *d*: 0.14, small effect size) and 30 minutes after exercise (Cohen’s *d*: 0.97, large effect size) in G2. Then again, no significant changes were observed in DAP (Figure 1).

**Figure 1:**
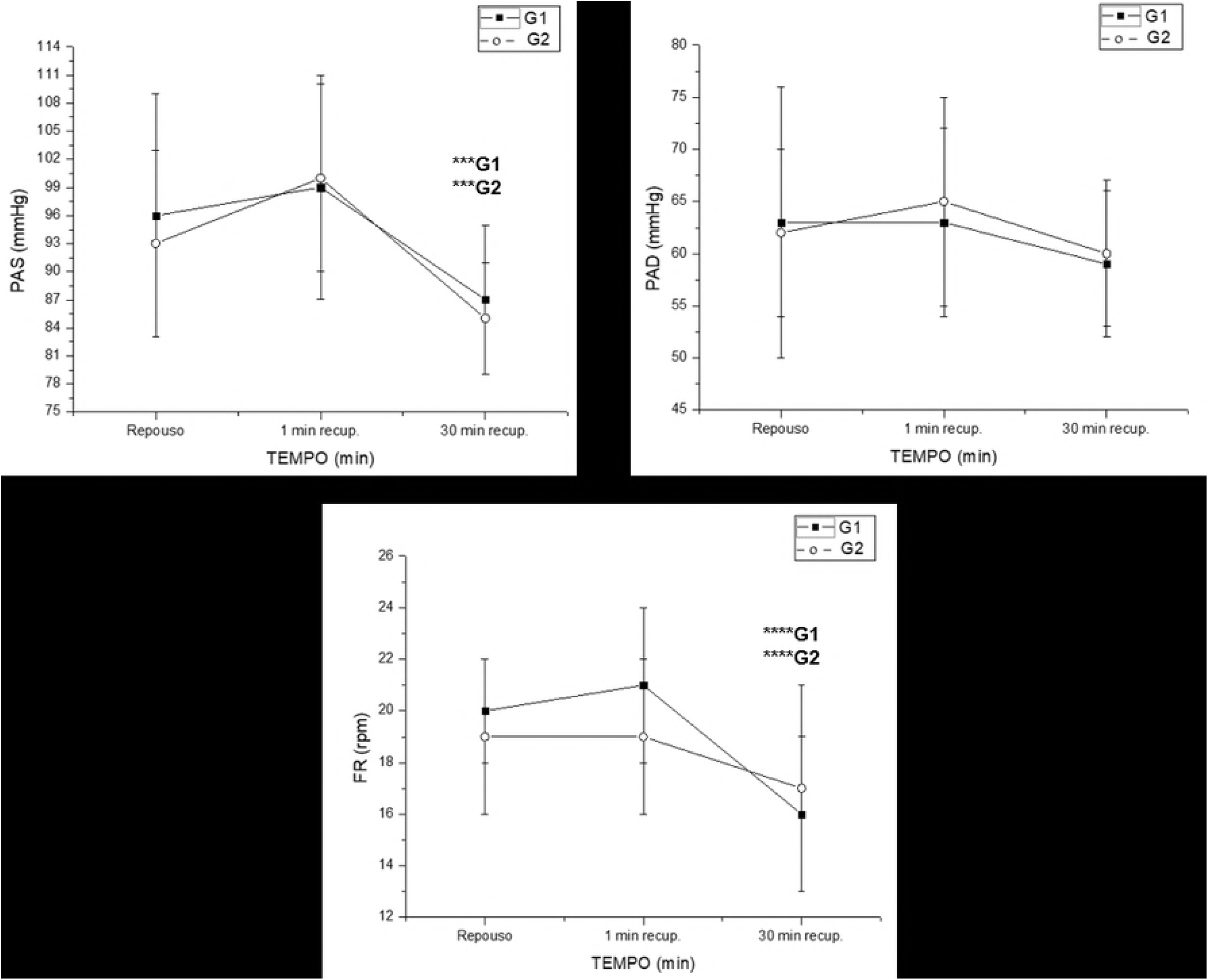
Mean values and respective standard deviations of SAP, DAP and f obtained at rest and during recovery from the submaximal aerobic exercise. ***G1: significant different in relation to rest in G1 (p<0.001); ***G2: significant different in relation to rest in G2 (p<0.001); ****G1: significant different in relation to rest in G1 (p<0.0001); mmHg: millimeters of mercury; cpm: cycles per minute;****G2: significant different in relation to rest in G2 (p<0.0001); SAP: systolic arterial pressure; DAP: diastolic arterial pressure; f: respiratory rate; G1: children with birth weight < 3400 grams; G2: children with birth weight > 3400 grams.

Figure 2 reveales HR, RR interval, HF and RMSSD deviate during recovery from exercise. HR was increased 0-5 minutes after exercise compared to rest in G2 (Cohen’s *d*: 1.52, large effect size). RR interval declined 0-5 minutes compared to rest in G1 (Cohen’s *d*: 0.7, medium effect size) and G2 (Cohen’s *d*: 1.56, large effect size). HF was reduced 0-5 minutes following exercise compared to rest in G2 (Cohens’*d*: 1.16, large effect size) and RMSSD was also reduced 0-5 minutes following exercise compared to rest in G2 (Cohens’*d*: 1.36, large effect size).

**Figure 2:**
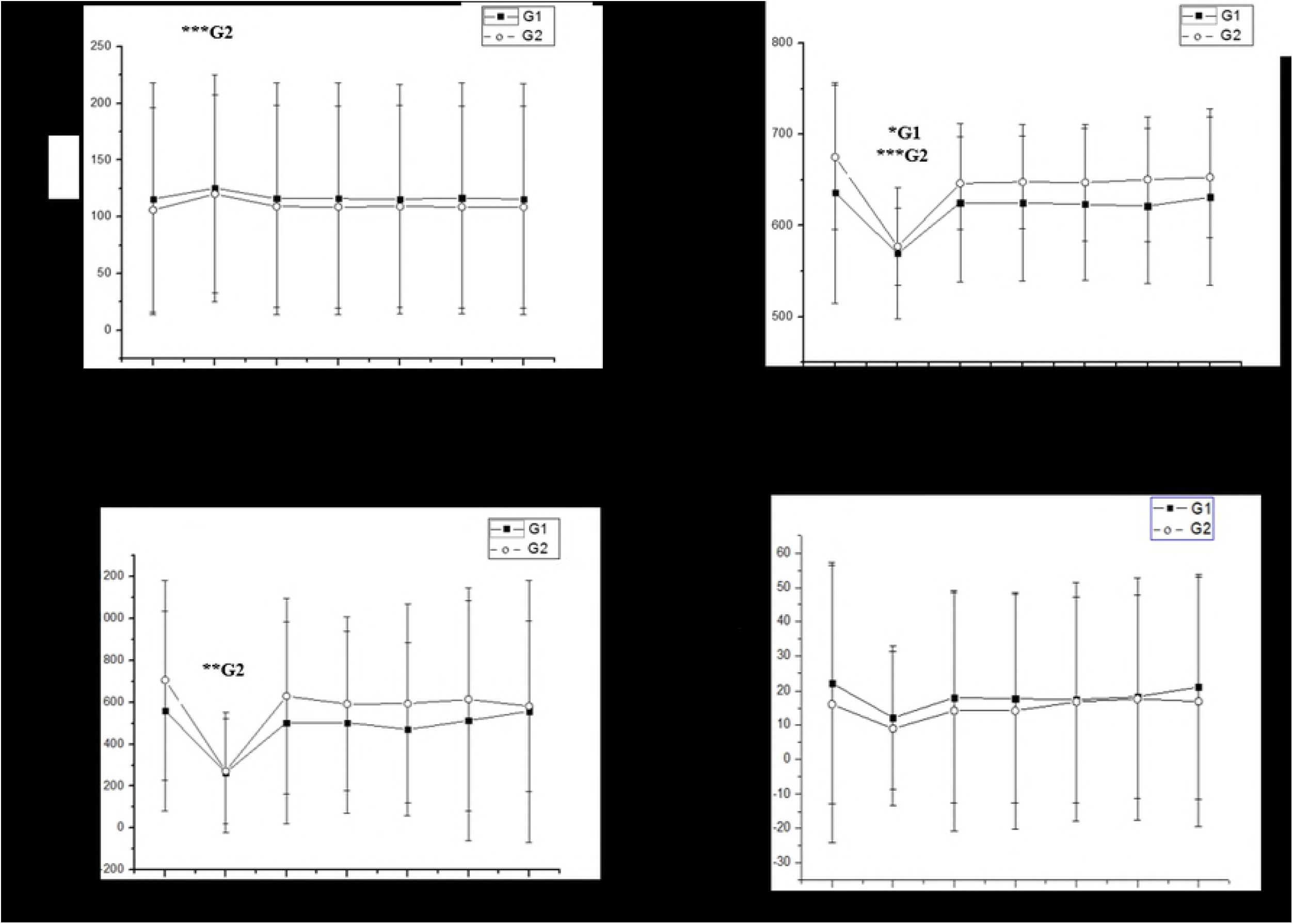
Mean values and respective standard deviations of HR, RR intervals, RMSSD and HF obtained at rest and during recovery from the submaximal aerobic exercise. *G1: significant different in relation to rest in G1 (p<0.05); **G2: significant different in relation to rest in G2 (p<0.01); ***G2: significant different in relation to rest in G2 (p<0.001); RMSSD: root-mean square of differences between adjacent normal RR intervals); HF: high frequency; ms: milliseconds; bpm: beats per minute; G1: children with birth weight < 3400 grams; G2: children with birth weight > 3400 grams.

In relation to symbolic HRV analysis, 0V increased 0-5 minutes after exercise compared to rest in G1 (Cohens’*d*: 1.01, large effect size) and G2 (Cohens’*d*: 1.31, large effect size), 2UV was decreased 0-5 minutes after exercise in G1 (Cohens’*d*: 1.31, large effect size) and G2 (Cohens’*d*: 0.9, large effect size) (Figure 3).

**Figure 3:**
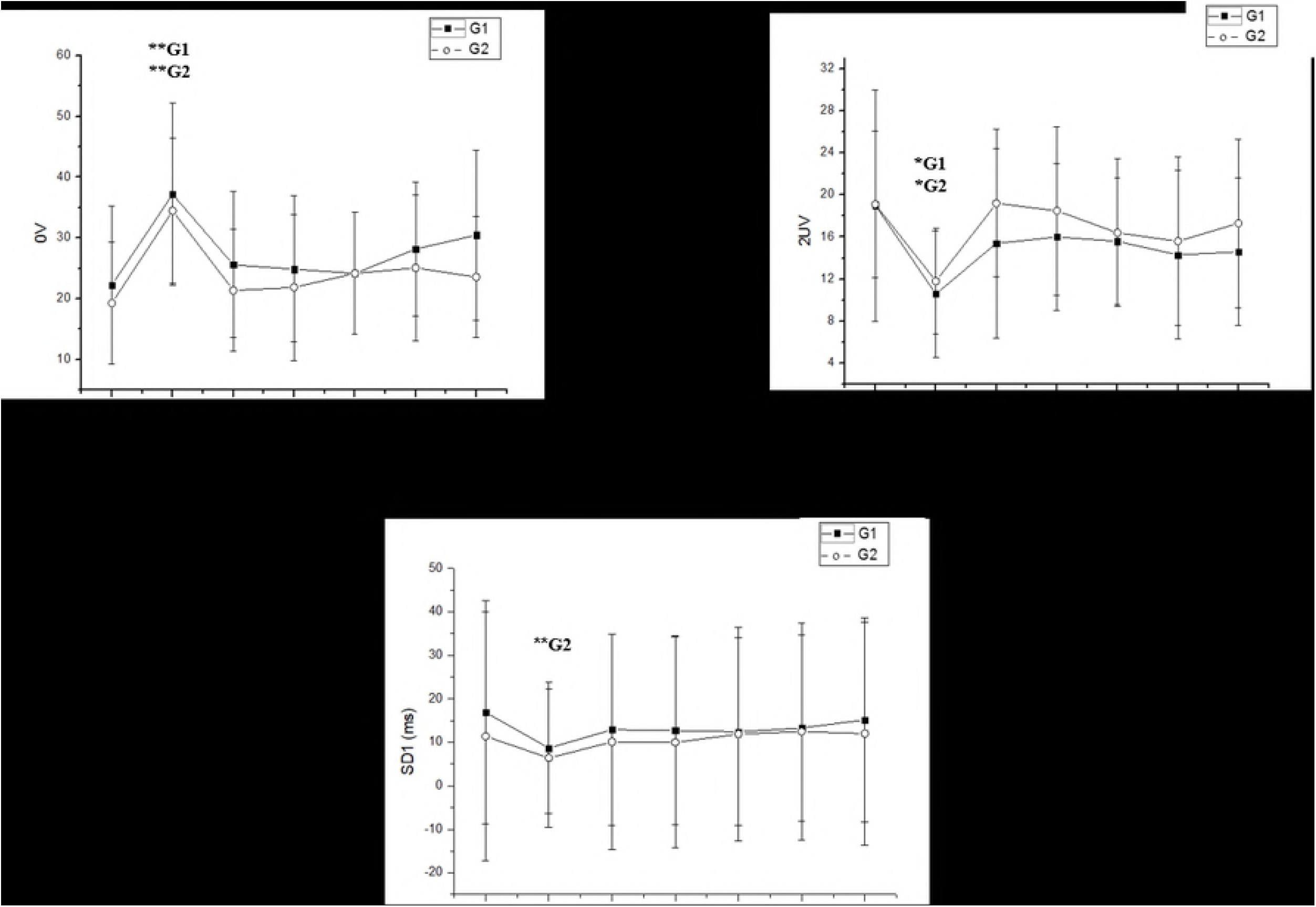
Mean values and respective standard deviations of SD1 and symbolic analysis obtained at rest and during recovery from the submaximal aerobic exercise. *G1: significant different in relation to rest in G1 (p<0.05); *G2: significant different in relation to rest in G2 (p<0.05); **G1: significant different in relation to rest in G1 (p<0.01); **G2: significant different in relation to rest in G2 (p<0.01); SD1: instantaneous recording of the variability of beat-to-beat ms: milliseconds; G1: children with birth weight < 3400 grams; G2: children with birth weight > 3400 grams.

Quantitative analysis of the Poincaré plot through SD1 index disclosed that it decreased 0-5 minutes after exercise compared to rest in G2 (Cohen’s *d*: 1.36, large effect size) (Figure 3).

Recurrence analysis was presented in Figure 4. Mean length was reduced 0-5 minutes after exercise compared to rest in G2 (Cohen’s *d*: 1.73, large effect size). Maximum length increased 0-5 minutes following exercise compared to rest (Cohen’s *d*: 1.57, large effect size) in G2. Recurrence rate increased 0-5 minutes after exercise compared to rest in G1 (Cohen’s *d*: 0.99, large effect size) and G2 (Cohen’s *d*: 0.99, large effect size). Determinism increased 0-5 minutes after exercise in G1 (Cohen’s *d*: 0.71, medium effect size) and G2 (Cohen’s *d*: 0.5, medium effect size). Laminarity increased 0-5 minutes after exercise compared to rest in G1 (Cohen’s *d*: 0.72, medium effect size) and G2 (Cohen’s *d*: 0.91, large effect size). Shannon Entropy enlarged 0-5 minutes after exercise compared to rest in G1 (Cohen’s *d*: 0.94, large effect size) and G2 (Cohen’s *d*: 0.76, medium effect size).

**Figure 4:**
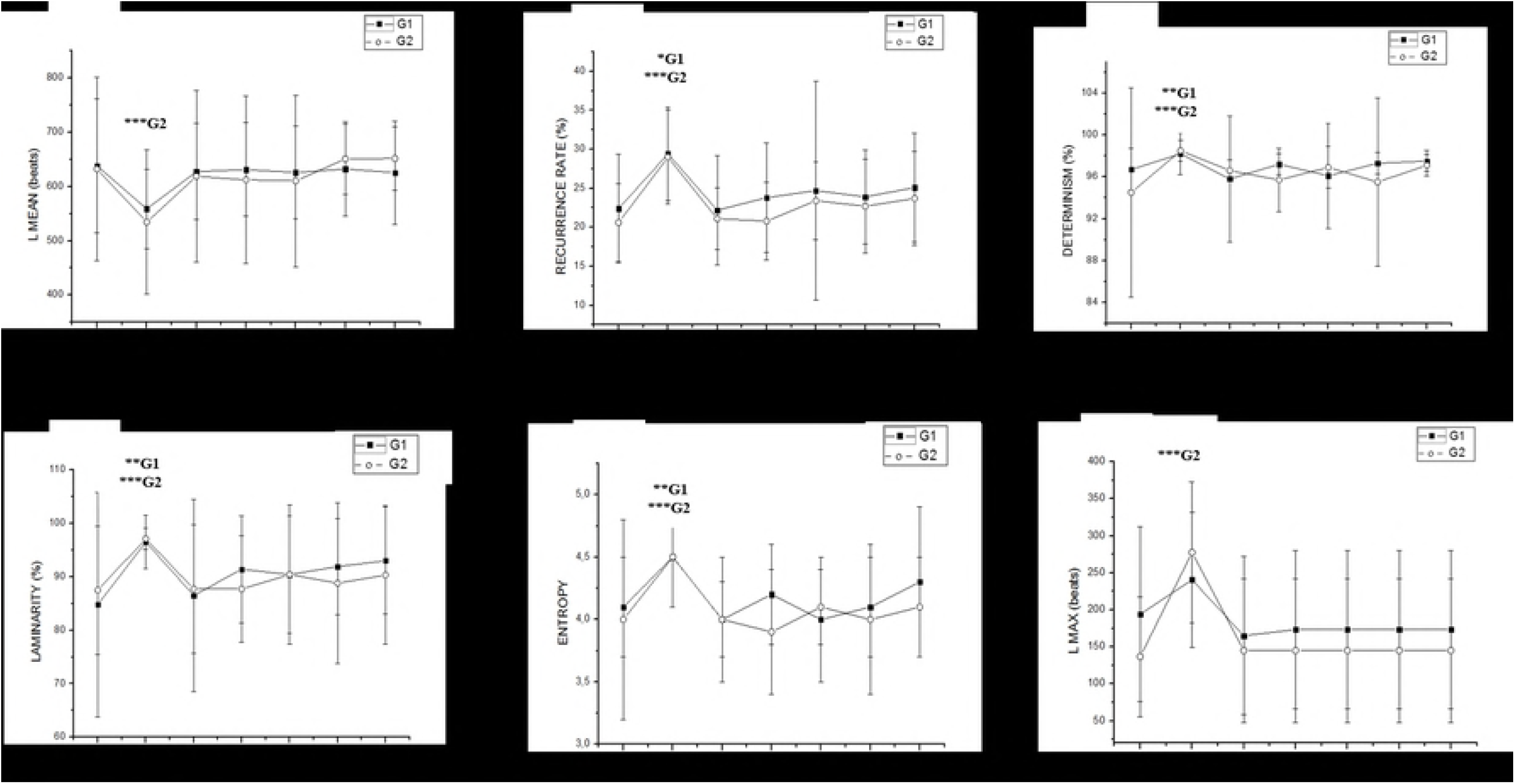
Mean values and respective standard deviations of recurrence analysis obtained at rest and during recovery from the submaximal aerobic exercise. *G1: significant different in relation to rest in G1 (p<0.05); *G1: significant different in relation to rest in G1 (p<0.01); ***G2: significant different in relation to rest in G2 (p<0.001); L MEAN: mean lenght; L MAX: maximum length; %: percentage; G1: children with birth weight < 3400 grams; G2: children with birth weight > 3400 grams.

Table 2 illustrates correlation between HRV and anthropometric variables. We documented weak correlation of fat percentage with 0V 0-5 minutes during recovery from exercise and recurrence rate 0-5 minutes during recovery from exercise. There was also weak correlation of BW with Determinism 5-10 minutes during recovery from exercise, Laminarity 5-10 minutes during recovery from exercise and Shannon Entropy 5-10 minutes during recovery from exercise (Table 2).

**Table 2:**
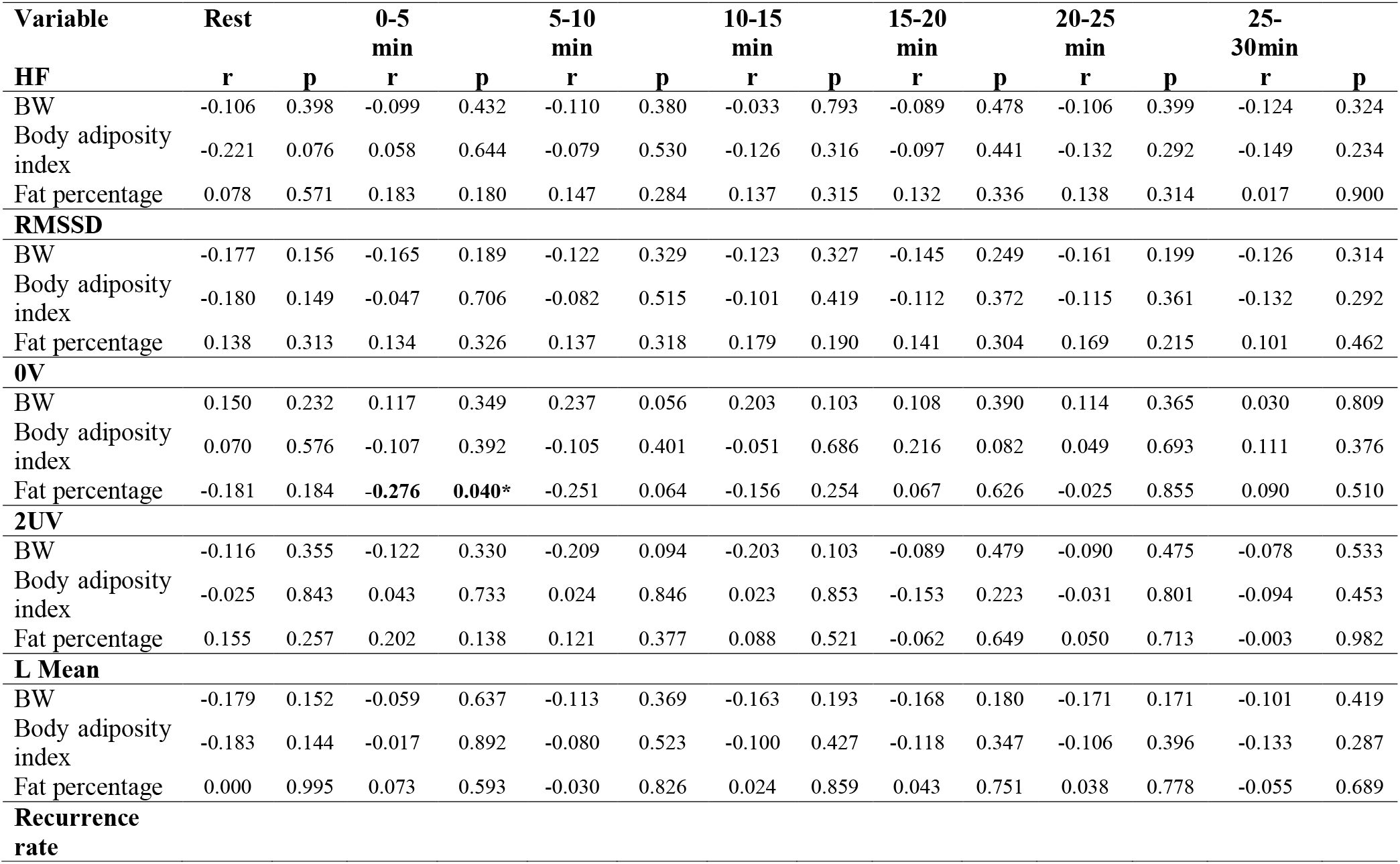
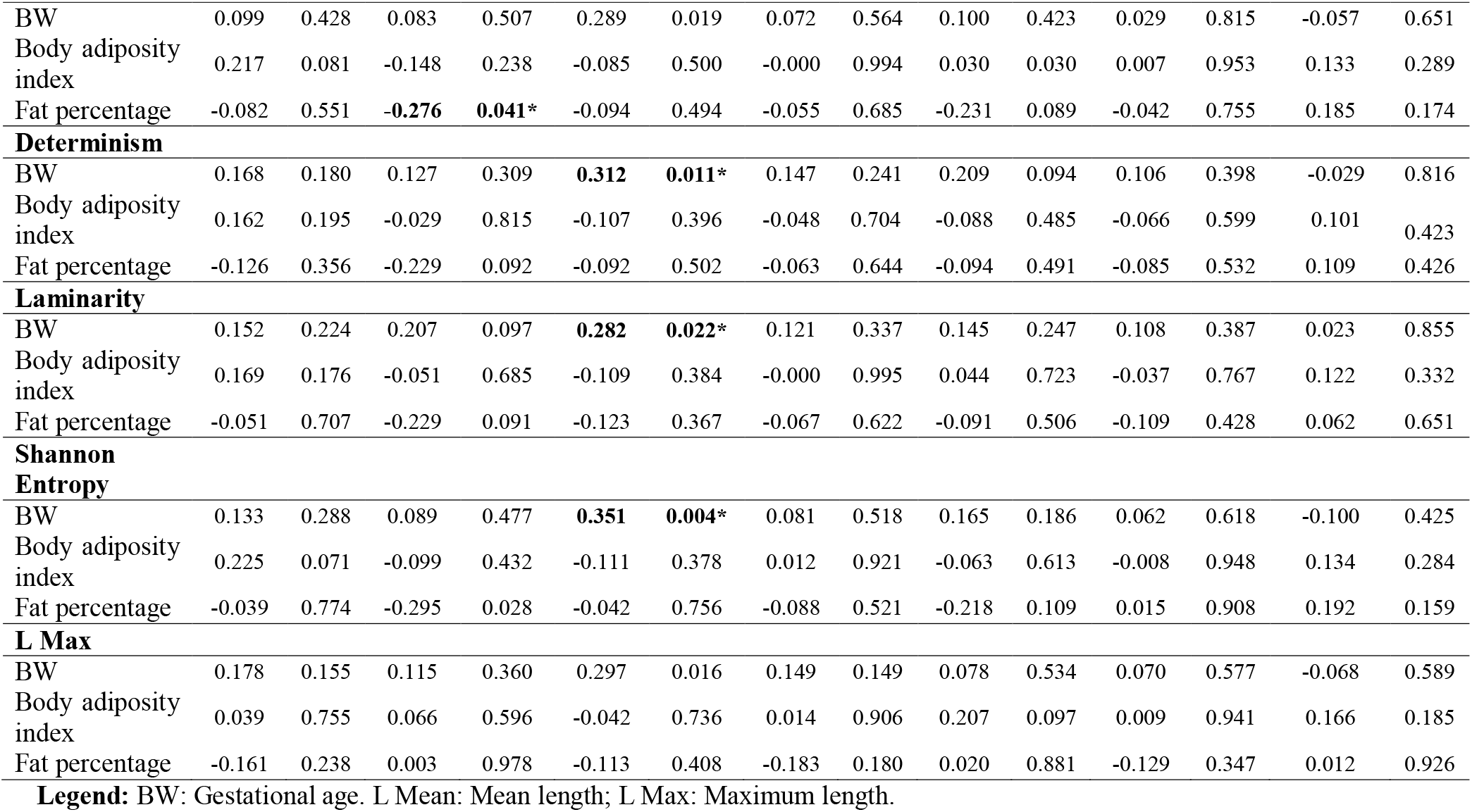
Correlation between anthropometric variables and HRV indices G1 in the moderate aerobic exercise protocol in. mmHg: millimeters of mercury. m: meters; kg: kilograms; mmHg: millimeters of mercury.

Linear regression established significant association between recurrence 0-5 minutes during recovery from exercise and fat percentage and between Shannon Shannon Entropy 5-10 minutes during recovery from exercise and BW (Table 3).

**Table 3.**
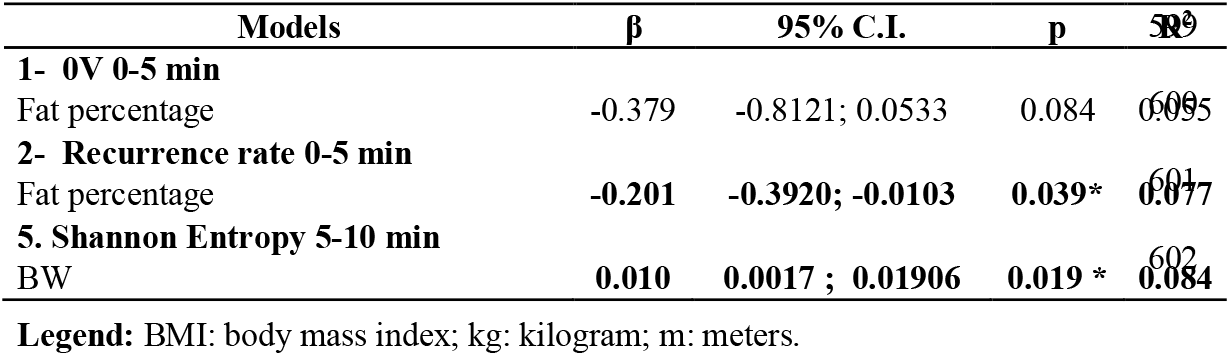
Linear regression between HRV and anthropometric variables.

## DISCUSSION

Childhood obesity can begin at any age, it occurs in a comparable way as in adults and is considered a global public health problem [42]. The etiology of obesity is multifactorial because it encompasses environmental, behavioral, organic, psychosocial and socioeconomic situations [43]. Thus, autonomic control of HR during physical effort is influenced by anthropometric factors by reason of the body composition and concentration of visceral fat deposits, which are strongly associated with cardiovascular diseases [44,45]. An earlier study indicated the possible impact of BW on ANS [19]. Yet, it is unclear if BW has significant impact in children during recovery from aerobic exercise, which is an important method to identify such autonomic changes [13].

In this way, our study was started to evaluate the involvement of BW and body composition on autonomic and cardiovascular recovery after submaximal aerobic exercise in children. As principal discoveries, we reported that: 1) f, SAP and DAP recovery were similar between children with higher and lower BW; 2) parasympathetic control of HR through linear HRV analysis was slightly delayed in children with higher BW; 3) nonlinear analysis of HRV indicated slower return following exercise in children with higher BW; 4) fat percentage and BW *slightly* influenced HRV recovery. It is imperative to mention that children with lower BW had increased fat percentage and body adiposity index.

The experimental conventions of this study were founded on HRV analysis during recovery from exercise. This procedure is often perfomed to detect cardiovascular diseases [14]. When related to acute submaximal aerobic physical exercise, it offers hemodynamic dysfunction that is occasionally unidentified at rest [13].

Based on our findings, SAP and f did not return to rest baseline levels 30 minutes after exercise. No significant changes in DAP were observed between before and during recovery from exercise. After physical exercise, SAP decreases due to peripheral vasodilatation [46], but, insufficient SAP decline may be indicative of cardiovascular disease and mortality [47]. Also, DAP normally remains stable during exercise [46]; and stimuli from the cerebral cortex effect a sudden increase in HR and f to the level that satisfies the requirements for metabolic gas exchange where, at the end of the exercise, they return to their baseline values [46].

We revealed that linear HRV analysis through power spectral density and time domain analysis demonstrated delayed return of HF band and RMSSD to rest values in children with higher BW. It designates that vagal control of HR offered slower return after exercise. This information is supported by previous studies [20–22,27] that had already evaluated the impact of BW on autonomic cardiac function. It was recognized that both low and high BW may be related to high risk of developing cardiovascular disease in both childhood and adulthood [48]. It was revealed that very low BW would be associated with immature autonomic activity [49], and that these individuals could exhibit reduced parasympathetic activity when attaining adulthood [21]. Nevertheless, the results remain inconclusive.

A previous study evaluated HR in 100 children aged 5-14 years old and demonstrated that there was a decrease in parasympathetic modulation in children with low BW, suggesting higher risk of developing cardiovascular and metabolic diseases during life in children with low BW. The aforesaid study strengthened the possibility that vagal withdrawal, rather than an increased sympathetic activity, may precede cardiovascular diseases in children with low BW [22].

Another study evaluated the ANS in 46 young adults aged 18-25 years old who consented to the handgrip test and it was observed that adults with low BW demonstrated an exaggerated increase in sympathetic response. The stated results support the hypothesis that adult subjects with low BW have adjustments in the autonomic modulation of HR [50].

In addition to linear HRV analysis, we performed nonlinear analysis. This is for the reason that the ANS is considered a complex, dynamic and non-linear system, since it is susceptible to numerous organic and environmental activities [51]. Symbolic and recurrence analysis of HRV are unalike the traditional linear analysis because it constructs the parameters based on nonlinear RR interval distribution [40].

Symbolic analysis of HRV exhibited that 0V and 2UV HRV parameters during recovery from exercise were similar in both groups. Previous studies involving pharmacological blockade and autonomic tests [52] indicated that the 0V% index epitomizes the cardiac sympathetic modulation and the 2ULV% index is related to cardiac vagal modulation.

Equally, recurrence HRV analysis evidenced that mean length was significantly decreased and maximum length was significantly increased 5 minutes after exercise in children with higher BW, whilst no significant change was recognized in the group with lower BW for those parameters. Recurrence rate, determinism, laminarity and Shannon Entropy were significantly increased 5 minutes after exercise compared to rest in both groups. The recurrence analysis is able to detect physiological changes [51] and non-stationary structural changes [53]. The lower the recurrence parameters values, the greater the chaotic response and more complexity in the system [51]. Our results propose that nonlinear HRV recovery from submaximal effort is delayed in children with higher BW.

Taken together, the research literature indicates that premature children and adults have greater likelihoods of developing cardiac autonomic dysfunction throughout life [22,50]. In contrast, our study evaluated term-born healthy children and separated them according to BW. With this in mind, we propose that higher BW may not have beneficial influences on the ANS.

We similarly examined the impact of body composition on the ANS. There was no significant change between higher and lower BW groups in relation to WC, abdominal circumference, WSR, WHR, BMI and conicity index. Nonetheless, we reported increased fat percentage and body adiposity index in children with reduced BW. This evidence raises the question: Which variable has greater influence on HRV recovery following exercise: BW, fat percentage or body adiposity index?

In order to resolve this question, we performed correlation and linear regression analysis. It was demonstrated that BW and current fat percentage has slight association with HRV recovery following exercise. Adjusted R^2^ revealed higher association of BW with nonlinear HRV recovery (BW: R^2^=0.084). In this way, we propose that BW has higher interaction with autonomic HR recovery.

Some points are worth highlighting from our study. Children of either gender between 9 and 11 years old were considered in this study. The children were not separated by gender because all children had homogeneous (pre-pubertal) maturational states. Guilkey *et al*. [27] evaluated the effects of autonomic modulation of the HR in children of the same age group and concluded that vagal activation during recovery from physical exercise was analogous amongst both boys and girls.

We evaluated healthy term-born children, henceforth, these results cannot be applied to populations with different ages and/or health status. We recognized that 21% of parents of children with lower BW and 28% of the parents of children with higher BW presented with cardiovascular disease. The research literature offered strong evidence regarding the influence of family history on risk factor for cardiovascular disease [54]. Yet, the groups were consistent regarding this variable.

## CONCLUSION

Term-born children with higher BW presented delayed autonomic recovery following aerobic submaximal exercise. BW and current fat percentage has impact on autonomic recovery from exercise. Our results draw attention to newborns with extreme high BW, since these outcomes provide important evidence that increased BW may be related to possible autonomic dysfunction and cardiovascular disorders in older ages. So, we highlight the importance of early detection of autonomic impairment before progression to the possible cardiovascular disorders.

## ACKNOWLEDGEMENTS

The authors would like to thank the IGeeks computer school, Prof. Anne Michelli Gomes and Prof. Letícia Santana de Oliveira for their technical assistance. The study received financial support from FAPESP (Number 2016/02994-1) and CNPq (Number 301079/2015-3).

## AUTHOR CONTRIBUTIONS

Juliana Edwiges Martinez Spada collected data, performed conduction of experiments, performed statistical analysis and draft the manuscript.

Fernando R. Oliveira performed conduction of experiments, statistical analysis and draft the manuscript.

David M. Garner reviewed statistical analysis, extensively reviewed the manuscript, English Grammar and Spelling.

Vitor E. Valenti supervised the study, wrote discussion section and gave final approval for the version submitted for publication.

All authors reviewed and approved the final version of the manuscript.

